# Postnatal development of somatosensory corticospinal projections in the mouse lumbar spinal cord

**DOI:** 10.64898/2026.02.27.708520

**Authors:** Antonia Maria Constantinescu, Lorenzo Fabrizi, Stephanie C. Koch

## Abstract

Corticospinal projections from the primary somatosensory cortex (S1) form a distinct descending pathway that engages spinal dorsal horn circuits and modulates sensory processing. Despite advances in our understanding of the role of this pathway in the adult, the postnatal maturation of somatosensory corticospinal projections remains poorly defined. Here, we provide a quantitative anatomical analysis of the postnatal development of corticospinal projections from the hindlimb representation of S1 (S1hl) to the lumbar dorsal horn in mice. Using retrograde tracing, we show that lumbar-projecting S1hl corticospinal neurons are first detected at postnatal day (P) 9 and reach adult-like numbers in S1hl by P12. Using anterograde tracing we then show that S1hl CST axonal projections are initially confined to the dorsolateral funiculus when they reach the lumbar cord at P9, but then rapidly invade the lumbar dorsal horn, reaching peak grey matter terminal density at P14. During this early innervation period, projections transiently extend beyond their mature termination zones in laminae III-V before retracting and becoming confined to superficial laminae I-II by P17. Together, these findings define three developmental phases of somatosensory corticospinal dorsal horn connections: an initial arrival phase, followed by grey matter ingrowth, and finally laminar refinement of the terminal projections. These results provide an anatomical framework for understanding how descending corticospinal somatosensory control becomes integrated into spinal circuits during the early postnatal period.

## Introduction

Precise sensorimotor behaviour depends on coordinated interactions between peripheral sensory input, local spinal circuitry, and descending supraspinal pathways. Among these pathways, the corticospinal tract (CST) is central to the refinement of spinal processing and motor output (Lemon and Griffiths, 2005). Although traditionally studied in the context of motor cortex, corticospinal projections from the primary somatosensory cortex (S1) form a distinct anatomical and functional subsystem. In adult rodents, corticospinal projections from S1 (S1-CST) terminate predominantly in the dorsal horn, where they co-localise with tactile afferent pathways and modulate sensory gain (Frezel *et al*., 2020; Moreno-Lopez *et al*., 2021; Guan *et al*., 2025; Wang *et al*., 2025). The termination patterns of these axonal projections have been suggested to define the behaviour they modulate, which implies S1-CST terminals within the spinal cord must target circuits with considerable accuracy as these behaviours are established in early life (Ueno *et al*., 2018).

The first three weeks of life in the rodent are marked by a maturation of sensorimotor reflexes and an extensive reorganisation of dorsal horn circuitry (Beggs *et al*., 2002; Fitzgerald, 2005). Anatomical studies of CST development have shown that corticospinal axons reach the spinal cord over this same time period as rodent behaviour becomes more refined (Donatelle, 1977; Schreyer and Jones, 1982; Gribnau et al., 1986; Joosten, Gribnau and Dederen, 1989; Curfs, Gribnau and Dederen, 1994; Martin, 2005), suggesting that descending circuits may shape behaviour through spinal circuit recruitment. However, to date, no studies have examined the maturation of S1-CST and its growth into and innervation of the lumbar spinal grey matter.

Here we provide a quantitative anatomical analysis of the postnatal maturation of corticospinal projections from the hindlimb representation of S1 (S1hl). Using complementary retrograde and anterograde tracing, we define the timing of corticospinal neuron arrival in the lumbar dorsal horn and quantify S1hl-CST spinal ingrowth, territory occupancy, and laminar targeting. By focussing upon the somatosensory corticospinal pathway, this study establishes an anatomical framework for future work on how early cortical activity engages spinal sensory circuits over the postnatal period.

## Materials and Methods

### Animals

All experiments were performed in compliance with the United Kingdom Animals (Scientific Procedures) Act of 1986. Animals had ad libitum access to water and food and were housed under 12-hour light/dark cycles.

Experiments were performed in wild type or homozygous *Emx1*^*Cre*^ C57Bl/6 mice of both sexes (n listed under each experiment). Retrograde tracing cohorts were wildtype mice analysed at P7, P9, P12, P14, P17, P19, P21 and adulthood (P33/P42 as indicated). Anterograde tracing cohorts were *Emx1*^*Cre*^ mice analysed at P7, P9, P12, P14, P17, P19 and adulthood (P42).

### Retrograde tracing from lumbar spinal cord

Retrograde tracing was performed by injection of Fluoro-Gold (FG) into lumbar dorsal horn at lumbar segment 4-5. FG was chosen for its lack of transsynaptic transport to selectively label the S1-CST lumbar projecting neurons (Catapano, Magavi and Macklis, 2002). Mice were placed in an induction chamber with 4% isoflurane in medical O_2_. Once induced, mice were positioned in a stereotaxic head frame (#51730U, Stoelting), and anaesthesia was maintained at 2% isoflurane in medical O_2_ via nosecone. Pups under the age of 15 days were induced in the stereotactic frame via nosecone and placed in custom neonatal head bars. The back was shaved and preoperative analgesic EMLA cream (AstraZeneca) was applied on the skin overlaying lumbar cord. Following disinfection with 4% Chlorhexidine (Hibiscrub), an incision was made in the skin and a laminectomy performed. To trace the developing corticospinal projections from the lumbar spinal cord, FG was injected in the lumbar cord (50–150 nl according to age, see Table 1) using a 1 µl Hamilton syringe (Model 7001 KH, Hamilton Company). The needle was left in place for 3 min after the injection to avoid leakage. Subsequently, the skin was sutured using Glueture (World Precision Instruments, US), animals received postoperative analgesia (Metacam, 2mg/kg) and were monitored and allowed to recover in a heated incubator. Brain and spinal cord tissue was processed for immunohistochemistry 5 days post injection, as outlined below.

**Table 1.** Volumes injected for retrograde/anterograde tracing of S1hl cortico-spinal neurons. FG injections were in the lumbar dorsal horn. AAV9 injections were in the S1hl.

| Age | Injected FG volume (nl) | Injected AAV9 volume (nl) |
| --- | --- | --- |
| P0 | - | 50 |
| P2 | 50 | 50 |
| P4 | 50 | 50 |
| P7 | 50 | 70 |
| P9 | 100 | 70 |
| P12 | 100 | 100 |
| P16 | 100 | 100 |
| P42 | 150 | 150 |

### Anterograde tracing of S1hl-CST projections

Mice were induced and placed within a stereotaxic frame as outlined above. The head was shaved and preoperative analgesic EMLA cream (AstraZeneca) was applied on the skin overlaying the skull. Following disinfection with 4% Chlorhexidine (Hibiscrub), an incision was made in the skin overlying the skull, and a craniotomy performed either via manual insertion of a 32G needle to pierce the skull and dura in pups, or via micro drill in adults (#58610, Stoelting). To trace the developing corticospinal projections from somatosensory cortex hindlimb region (S1hl), a Cre-dependent anterograde viral tracer (AAV9 hSyn-DIO-hM4D (G_i_)-mCherry, #44362-AAV9, Addgene; working titre ≥ 1×10^13^ vg/mL, 50-150 nl according to age, see Table 1) was injected in S1hl using a 1 µl Hamilton syringe (Model 7001 KH, Hamilton Company) attached to a manual microinjector (Model 5000, Kopf). The needle was left in place for 3 min after injection to avoid leakage. Subsequently, the skin was sutured using Glueture (World Precision Instruments, US), animals were given postoperative analgesia (Metacam, 2mg/kg) and were monitored and allowed to recover in a heated incubator. Brain and spinal cord tissue was processed for immunohistochemistry 7 days post injection, as outlined below. Age-specific S1hl stereotaxic coordinates were determined using Nissl cytoarchitectonic landmarks and atlas alignment (Supp. Fig. 1; Table 2). S1hl and M1 were differentiated according to layer 4 and layer 5 thickness and cell body orientation (Yamawaki, 2014). Injection site placement and injectate spread was verified histologically for each animal as outlined below.

**Table 2.** Stereotaxic coordinates for S1hl cortical injections. AP (antero-posterior location from the λ landmark), ML (medio-lateral location from the midline), and DV (dorso-ventral, or depth of injection).

| Age | AP from $\lambda$<br>(mm) | ML from<br>midline (mm) | DV from skull<br>surface (mm) |
| --- | --- | --- | --- |
| P0 | 1.3 | 1.1 | 0.5 |
| P3 | 1.7 | 1.2 | 0.5 |
| P5 | 2.4 | 1.6 | 0.6 |
| P7 | 2.6 | 1.6 | 0.7 |
| P9 | 2.6 | 1.6 | 0.8 |
| P10 | 3 | 1.6 | 0.8 |
| P14 | 3.8 | 1.8 | 0.9 |
| P16 | 4 | 1.9 | 0.9 |
| P42 | 4.1 | 1.5 | 1 |

### Tissue processing and immunohistochemistry

Five days post spinal FG injection, or 7 days following cortical AAV9 hSyn-DIO-hM4D (Gi)-mCherry, mice were transcardially perfused with 0.9% saline solution with heparin followed by ice-cold 4% paraformaldehyde (PFA) in PBS prior to spinal cord (lumbar levels L4 and L5) and brain dissection. The tissue was post-fixed for 2 hours at 4°C in 4% PFA. Tissues were incubated in 30% sucrose/PBS for 3 nights, or until completely immersed in sucrose and subsequently sectioned at 40μm on a LEICA SM 2000 R microtome. Prior to immunohistochemical staining, sections were permeated for 30 minutes in 50% ethanol solution then washed 3 times for 10 min in PB. These were then blocked for 1h in 10% donkey serum and 10% Triton X solution in PBS. Sections were then incubated over three nights with primary antibodies at 4°C (see Table 3 for details). Primary antibody staining was detected with fluorophore–conjugated secondary antibodies (1:300; Jackson Laboratories). When DAPI co-stain was used, staining was performed in the last wash, after secondary antibody incubation, for 10 minutes at a concentration of 1:10000. 40 µm coronal brain sections were mounted onto gelatinised slides.

**Table 3.** List of primary and secondary antibodies used in immunohistochemistry.

| Antibody | Host species | Concentration | Supplier | Protocol |
| --- | --- | --- | --- | --- |
| Primary |  |  |  |  |
| Ds-Red | Rabbit | 1:500 | Takara, 632496 | Direct |
| vGluT1 | Guinea Pig | 1:1000 | Millipore, AB5905 | Direct |
| PKCγ | Rabbit | 1:10000 | Santa Cruz Biotech, D0604 | TSA |
| NeuN | Mouse | 1:2000 | Millipore, MAB377 | Direct |
| Fluoro-Gold | Rabbit | 1:500 | Millipore, AB153-1 | Direct |
| Secondary |  |  |  |  |
| Cy2 anti rabbit | Donkey | 1:300 | Jackson, 711545152 | Direct |
| Cy3 anti rabbit | Donkey | 1:300 | Jackson, 711225152 | Direct |
| Cy2 anti guinea pig | Donkey | 1:300 | Jackson, 706165148 | Direct |
| Cy5 anti guinea pig | Donkey | 1:300 | Jackson, 706175148 | Direct |

Nissl staining was used in a subset of sections at each age to identify S1hl and associated layers. To do so, slides were dipped in (1) distilled water (1 second) and then submerged in (2) 0.1 cresyl violet (30 seconds). Following this, the slides were then dipped for 20 seconds in the following solutions: (3) distilled water, (4) 50% ethanol, (5) 70% ethanol, (6) 95% ethanol, (7) xylene, (8) xylene. Slides were then coverslipped using DPX mounting medium.

### Imaging

Nissl-stained sections images were captured using a Zeiss Axio Scan Z1 slide scanner microscope (Zeiss, Germany).

Cortical and spinal sections for retrograde studies were imaged using a LEICA DM R fluorescent microscope using x1.6, x5, x10, and x20 objectives and analysed using Fiji/ImageJ software.

Cortical and spinal sections for anterograde S1hl-CST studies were imaged with an inverted confocal microscope (SoRa Yokogawa CSU-W1 series, Nikon) and analysed using Fiji/ImageJ software.

### Cell counting (retrograde tracing)

FG^+^ neurons in S1hl layer 5 contralateral to the spinal injection site were counted manually using the Fiji Cell Counter plugin within a fixed region of interest (ROI) of 0.49 mm^2^. S1hl was identified in these sections using a combination of the Allen Brain Atlas and reference sections from Nissl staining (detail above; Supp. Fig. 1). Cell counts were expressed as the average number of FG labelled cells in n=5 sections per animal (n=3 animals per age other than P7 and P12 where n=2). DAPI^+^ nuclei within the same ROI were automatically quantified in Fiji/ImageJ by applying a fluorescence threshold to generate a binary mask, followed by automated cell counting using the Analyze Particles plugin. DAPI^+^ counts in layer 5 of S1hl were not significantly different across ages (Supp. Fig. 2).

### Quantification of dorsal horn occupancy (anterograde tracing)

S1hl-CST occupancy was quantified as the proportion of the contralateral lumbar dorsal horn area occupied by mCherry signal (thresholded at mean + 2 SD of the fluorescence across the entire section), as measured within Fiji/ImageJ. Quantification was performed on n=3 sections per animal (n=3 animals per age) across lumbar segments L3–L5. The distribution and density of mCherry signal (S1hl-CST occupancy) within the dorsal horn was visualised using the ‘spline 16’ interpolation method of the matplotlib library and Python script kindly provided by the Cordero-Erausquin lab (Moreno-Lopez et al., 2021).

### Laminar distribution analysis

Laminar distribution of S1hl-CST projections within the dorsal horn was quantified as the proportional overlap of mCherry (fluorescence thresholded at mean + 2 SD of the fluorescence across the entire section) with laminar markers (PKCγ for lamina II_i_; vGluT1 for laminae III–V) (Malmberg *et al*., 1997; Varoqui *et al*., 2002). Due to the absence of layer specific markers for superficial laminae, mCherry distribution within laminae I-II_o_ was estimated as (100% - (PKCγ% overlap + vGluT1% overlap)).

### Statistics

Developmental trajectories for FG^+^ counts, S1hl-CST dorsal horn occupancy and laminar distribution were modelled with polynomial regressions. To determine the optimal polynomial degree, models from constant to third order polynomial were fitted, and the corresponding R^2^ values were plotted. The model degree selected corresponded to the lowest degree at which the R^2^ curve began to plateau. To estimate the age at which FG^+^ counts, S1hl-CST dorsal horn occupancy and laminar distribution reached adult-like maturity, a model-based 95% confidence interval overlap approach was used. Maturity was defined as the earliest age at which the 95% confidence interval (CI) of the polynomial regression modelling the developmental trajectories of FG^+^ counts, occupancy and laminar distribution fully contained the adult group 95% CI. Data are presented as mean ± 95% CI.

DAPI^+^ counts and S1hl mCherry spread are presented as mean ± standard deviation (SD) and analysed using a one-way ANOVA.

## Results

### Developmental emergence of lumbar-projecting S1hl-CST neurons

We first sought to determine when CST projections from the hind limb region of the primary somatosensory cortex (S1hl-CST) reach the lumbar dorsal spinal cord using retrograde labelling of S1hl-CST projections. Retrograde FG injections into lumbar dorsal horn (Fig. 1A-B) yielded very few, weakly labelled S1hl L5 neurons (FG^+^) at P7, while clearly labelled neurons were visible at P9 and older ages (Fig. 1C). Labelled FG^+^ cells were found to co-express NeuN, indicating a neuronal phenotype and specificity of labelling (Fig. 1D). To determine the age at which the number of FG^+^ neurons in S1hl reached adult levels, we modelled the developmental trajectory of FG^+^ cell counts using a second-order polynomial and compared the resulting 95% confidence interval with that of adults (P33: 0.02 ± 0.002). FG^+^ cell count reached adult levels at P12 (95% CI overlapped fully with that of adults, Fig. 1E, black cross), indicating that the number of lumbar-projecting S1hl corticospinal neurons stabilises by P12 (Fig. 1E).

**Fig 1.**
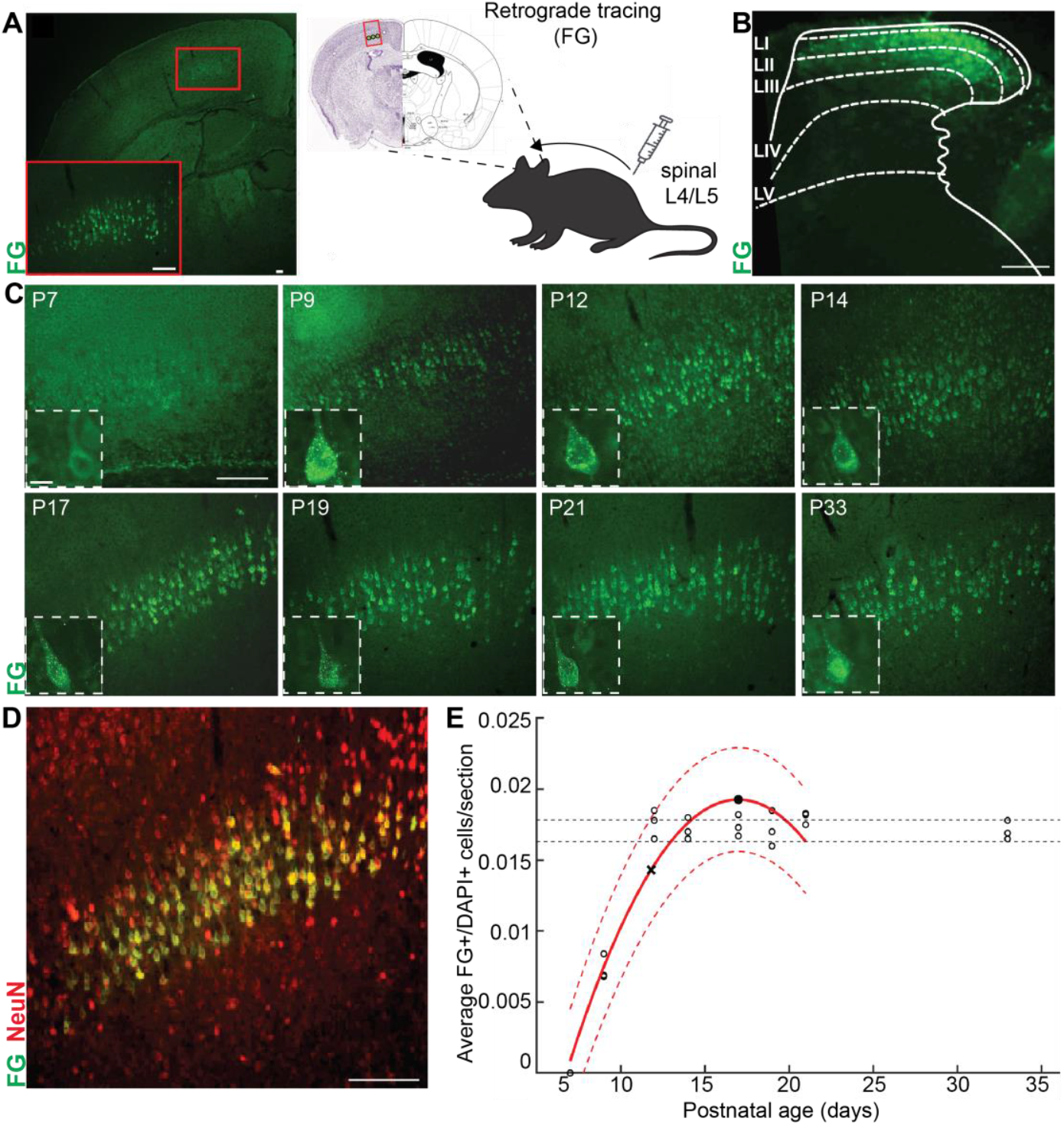
S1hl-CST lumbar projections do not reach the spinal cord before P7. **(A)** (left) Representative coronal section showing FG-labelled corticospinal neurons (FG^+^) in the S1 hindlimb (S1hl) cortex of a P21 mouse following retrograde tracer injection at spinal level L4– L5. Red boxes indicate higher magnification within S1hl L5. (right) Schematic illustrates the injection site and retrograde transport of FG to cortical projection neurons. **(B)** Representative image from a P21 animal showing FG spinal injection site in L4. White outlines indicate dorsal horn laminae. **(C)** Representative FG^+^ (green) corticospinal neurons in S1hl across postnatal ages P7, P9, P12, P14, P17, P19, P21, P33. Insets (white dashed boxes) show FG^+^ neurons at each age. **(D)** Representative cortical section at P21 showing co-labelling of FG (green) with the neuronal marker NeuN (red). Overlap shown in yellow. **(E)** Quantification of retrogradely labelled FG^+^ neurons in the cortex normalised to DAPI. Each data point represents a single animal. The solid line represents a 2^nd^ order polynomial fit to the data and dashed lines the boundaries of the 95% CI. Horizontal dotted lines represent the boundaries of the 95% CI in adults. Black cross represents the first postnatal day from when there was no more significant difference in the number of FG^+^ CSNs compared to adults. Scale bars: (A, B, D) 100µm, C 100 µm for larger panels and 10 µm for insets.

These data demonstrate that lumbar-projecting S1hl corticospinal neurons are first detectable at P9 and reach adult-like numbers by P12.

### Developmental innervation and pruning of S1hl-CST projections in the lumbar dorsal horn grey matter

Having determined that S1hl-CST projections reach the lumbar cord after P7, we next examined the timing and specificity of their innervation into the lumbar dorsal horn grey matter. To do this, we anterogradely labelled S1hl-CST projections by viral injection into the somatosensory cortex and analysed axonal projections within the grey matter (Fig. 2A).

**Fig 2.**
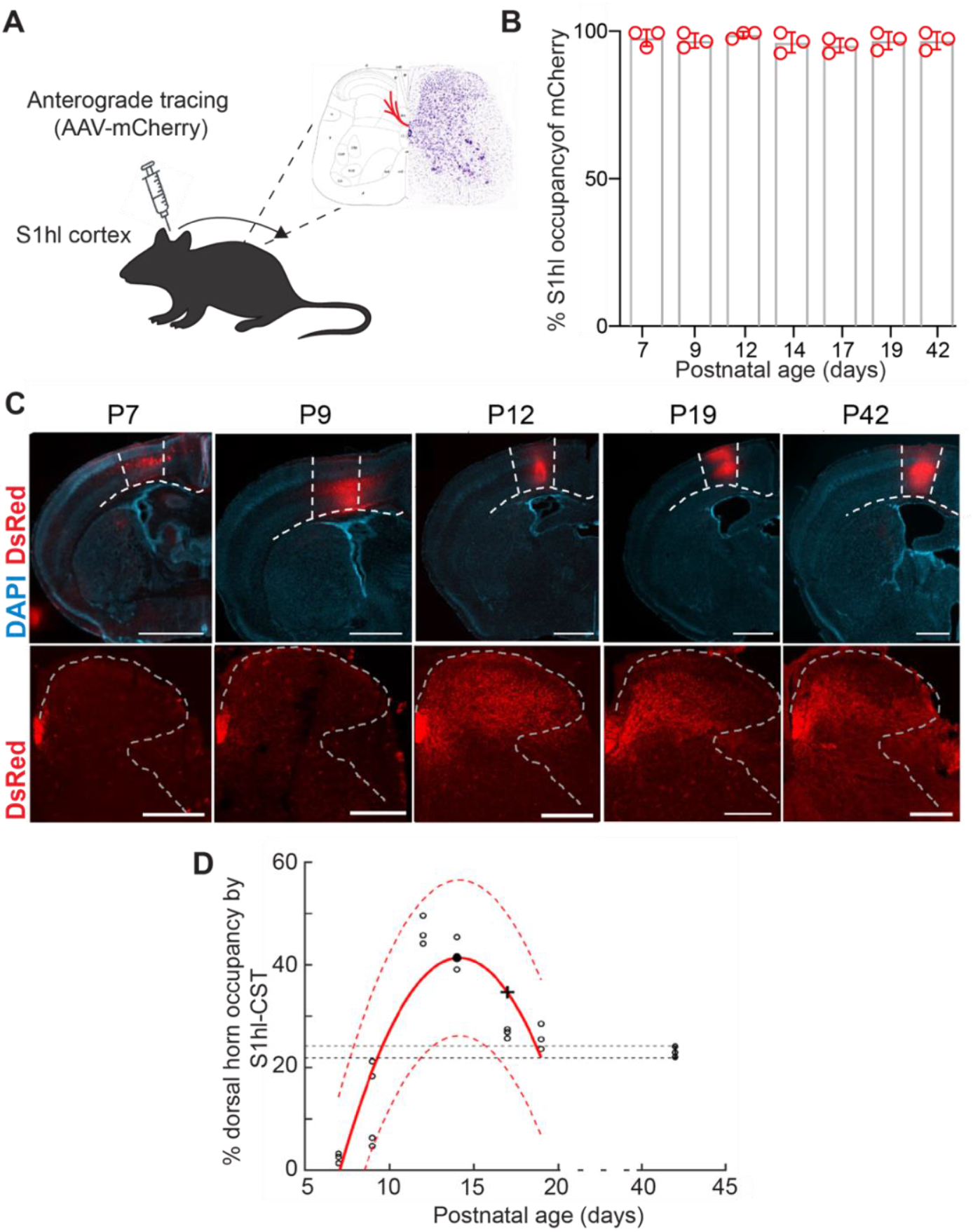
S1hl-CST axonal grey matter occupancy follows a developmental biphasic maturation of extension followed by retraction. **(A)** Schematic illustrates the injection site and anterograde transport of mCherry to lumbar spinal cord. **(B)** mCherry injectate spread within S1hl L5 at P7, P9, P12, P14, P17, P19 and P42 (mean ± SD; n=3 animals/age). **(C)** Representative coronal brain (upper) and lumbar (L3-5) spinal cord (lower) sections showing mCherry expression. White dashed lines represent S1hl boundaries and spinal cord outline. Scale bar 1000µm. **(D)** Percentage of dorsal horn S1hl-CST occupancy at different ages. Each circle represents a single animal. The solid line represents a 2^nd^ order polynomial fit to the data and dashed lines the boundaries of the 95% CI. Horizontal dotted lines represent the boundaries of the 95% CI in adults. Full black circle marks the peak S1hl-CST occupancy. Black cross represents the first postnatal day from when there was no more significant difference in the grey matter occupancy compared to adults. Scale bars: 1000 µm.

Successful viral targeting of S1hl was confirmed by inspection of mCherry expression within S1hl layer 5, the output layer of S1hl-CST projections. mCherry was expressed in 98.33% ± 2.89% of L5 of S1hl at P7; 97.33% ± 2.52% at P9; 99.33% ± 1.16% at P12; 96.67% ± 3.51% at P14; 95.67% ± 2.52% at P17; 97.33% ± 3.06% at P19 and 97.33% ± 3.06% at P42; Fig 2B, one-way ANOVA, n.s.). mCherry was expressed in the dorsal funiculus of the lumbar segment at all ages, suggesting that the timing of viral transfection was sufficient for mCherry expression and that the viral injections were targeted to the S1hl-CST (Fig. 2C).

In agreement with our retrograde tracing, S1hl-CST spinal dorsal horn axonal innervation was minimal at P7, with fluorescence found to be largely restricted to the dorsolateral funiculus (Fig. 2C-D). From P9 onwards, innervation and branching increased rapidly throughout the dorsal horn, reaching peak grey matter occupancy at P14 (mean ± SD = 42.06 ± 3.14% of dorsal horn territory; Fig. 2D).

To determine the age at which S1hl-CST axonal branching reaches maturity, we modelled the developmental trajectory of S1hl-CST dorsal horn occupancy with a second-order polynomial and compared its 95% confidence interval with those in adult (P42) (23.05 ± 1.03%). Occupancy at P14 was significantly higher than in adults (CI did not overlap), but by P17 (26.69 ± 0.92%) 95% CI overlapped fully with that of adults (Fig. 2D, black cross), indicating a developmentally mature pattern of axonal branching. This biphasic profile of extension and retraction suggests an exuberant phase of grey matter innervation in the second postnatal week followed by refinement during the third.

### Transient superficial spinal dorsal overextension and later refinement of S1hl-CST projections

Having established a biphasic developmental profile of S1-CST axonal innervation into the dorsal horn, we next determined the laminar distribution of these early aberrant projections. To do so, we quantified the developmental distribution of S1hl-CST projections among dorsal horn laminae by their overlap with laminar markers PKCγ in lamina (L) II_i_, and vGluT1 in LIII-V (Abraira and Ginty, 2013; Fig. 3A-B).

**Fig 3.**
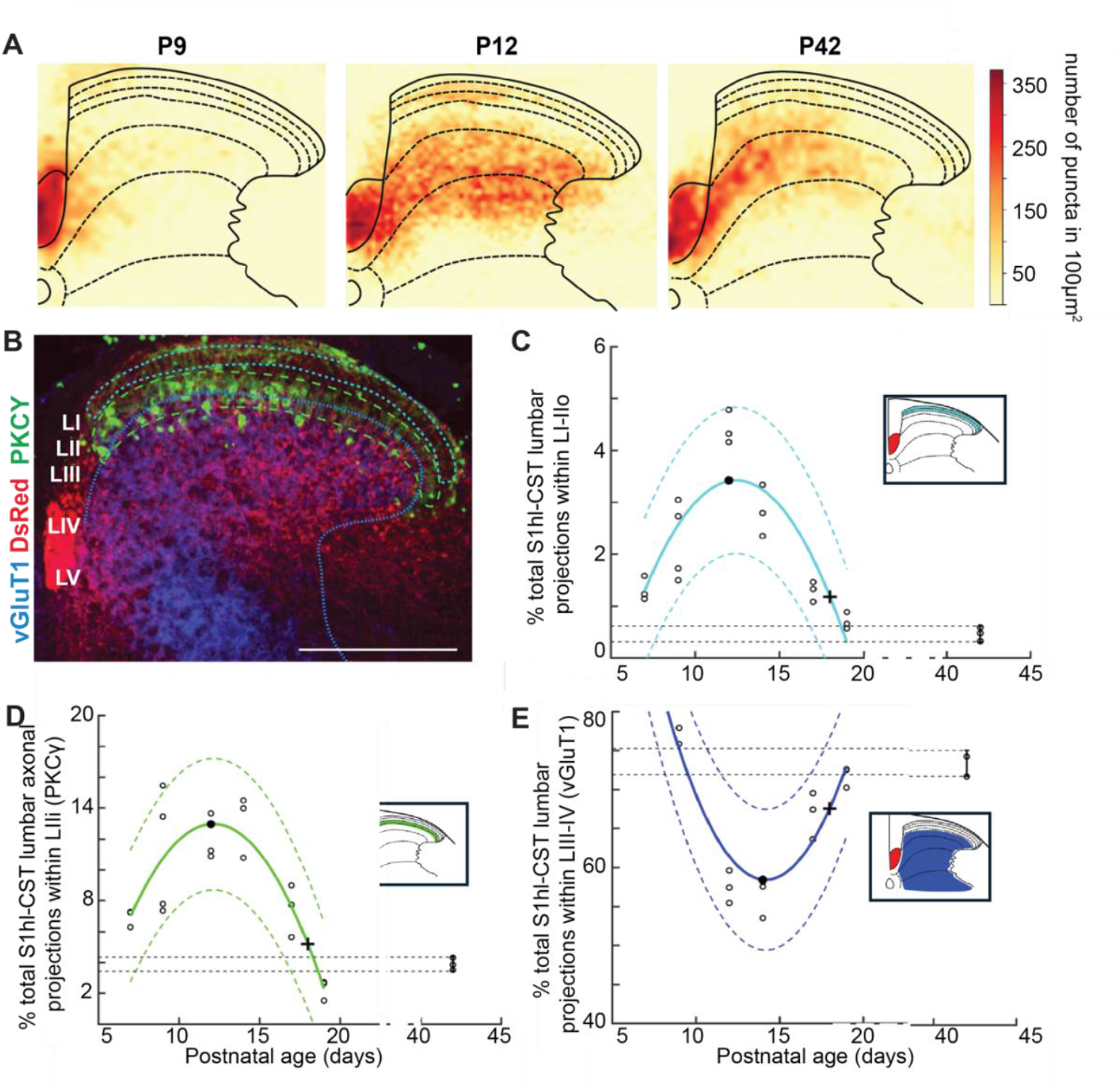
S1hl-CST lumbar projections extend to superficial laminae during the first two postnatal weeks and retract to deeper laminae over the third postnatal week. **(A)** Heatmaps of S1hl-CST grey matter projections at P9, P12 and P42 (n=4 animals/age). Heatmap colour intensity represents the number of mCherry^+^ puncta. **(B)** Representative P12 transverse L4 spinal section immunostained with DsRed (red), PKCγ (green), and vGluT1 (blue). Dashed lines represent laminar boundaries. **(C)** Percentage of total S1hl-CST axonal projections within LI/II_o_, **(D)** LII_i_ and **(E)** LIII-V across postnatal development. Total of C-E will total 100% for each animal. Each circle represents a single animal. The solid lines represent a 2^nd^ order polynomial fit to the data and dashed lines the boundaries of the 95% CI. Full black circles mark the peak S1hl-CST percentage occupancy for each laminar compartment. Black cross represents the first postnatal day from when there was no more significant difference compared to adults. Scale bars 500 µm.

Contrary to innervation patterns seen in the adult (Liu *et al*., 2018; Frezel *et al*., 2020), S1hl-CST axonal projections were found to transiently extend beyond LIII-IV and into the superficial dorsal horn in the early postnatal period. S1hl-CST projections displayed a biphasic innervation into superficial LI-II_o_ and II_i_, showing initial extension, followed by refinement (P7: LI-LII_o_: 1.32 ± 0.19, LII_i_: 8.79 ± 0.88; P9: LI-LII_o_: 2.43 ± 0.67, LII_i_:14.45 ± 4.61; P12: LI-II_o_: 4.753 ± 0.4206%, LII_i_: 14.55 ± 1.840%, ; P14: LI-LII_o_: 2.83 ± 0.4, LII_i_: 15.96 ± 2; P17: LI-LII_o_:1.29 ± 0.16, LII_i_:7.45 ± 1.39; P19: LI-LII_o_: 0.4667 ± 0.1305%, LII_i_: 3.893 ± 0.3911%; P42: LI-LII_o_: 0.47 ± 0.11, LII_i_: 3.89 ± 0.32; Fig. 3C-D). An inverse relationship was instead seen with regards to deep dorsal horn innervation (LIII-V - P7: 87.42 ± 1.18; P9: 79.65 ± 3.99; P12: 57.52 ± 1.71; P14: 56.43 ±2.05; P17 66.88 ± 2.44; P19: 71.82 ± 1.11; P42: 73.61 ± 1.2, Fig. 3E). These data indicate that S1hl-CST axonal laminar specificity within the lumbar dorsal horn emerges through a transient overextension into superficial layers that is refined to an adult-like pattern by the end of the third postnatal week.

### Emergence of putative presynaptic terminals in the developing S1hl-CST

Since anterograde tracing labels both synaptic terminals and fibres of passage, we aimed to determine whether initial S1hl-CST projections show evidence of synaptic contacts within the lumbar dorsal horn. To label presynaptic terminals of S1hl-CST projections, we co-stained spinal sections of anterogradely traced axons with vGluT1, which labels terminals of both myelinated afferents and CST projections (Alvarez *et al*., 2004; Broadhead *et al*., 2020). Colocalisation of mCherry and vGluT1 was detectable in medial deep dorsal horn CST termination zone as early as P9 (Fig. 4A-D). This colocalisation persisted at P12 and into adulthood (P42), consistent with early acquisition of presynaptic features. While colocalisation does not indicate functionality, these anatomical data suggest that descending somatosensory corticospinal projections form putative synaptic contacts within the dorsal horn shortly after entering spinal grey matter.

**Fig 4.**
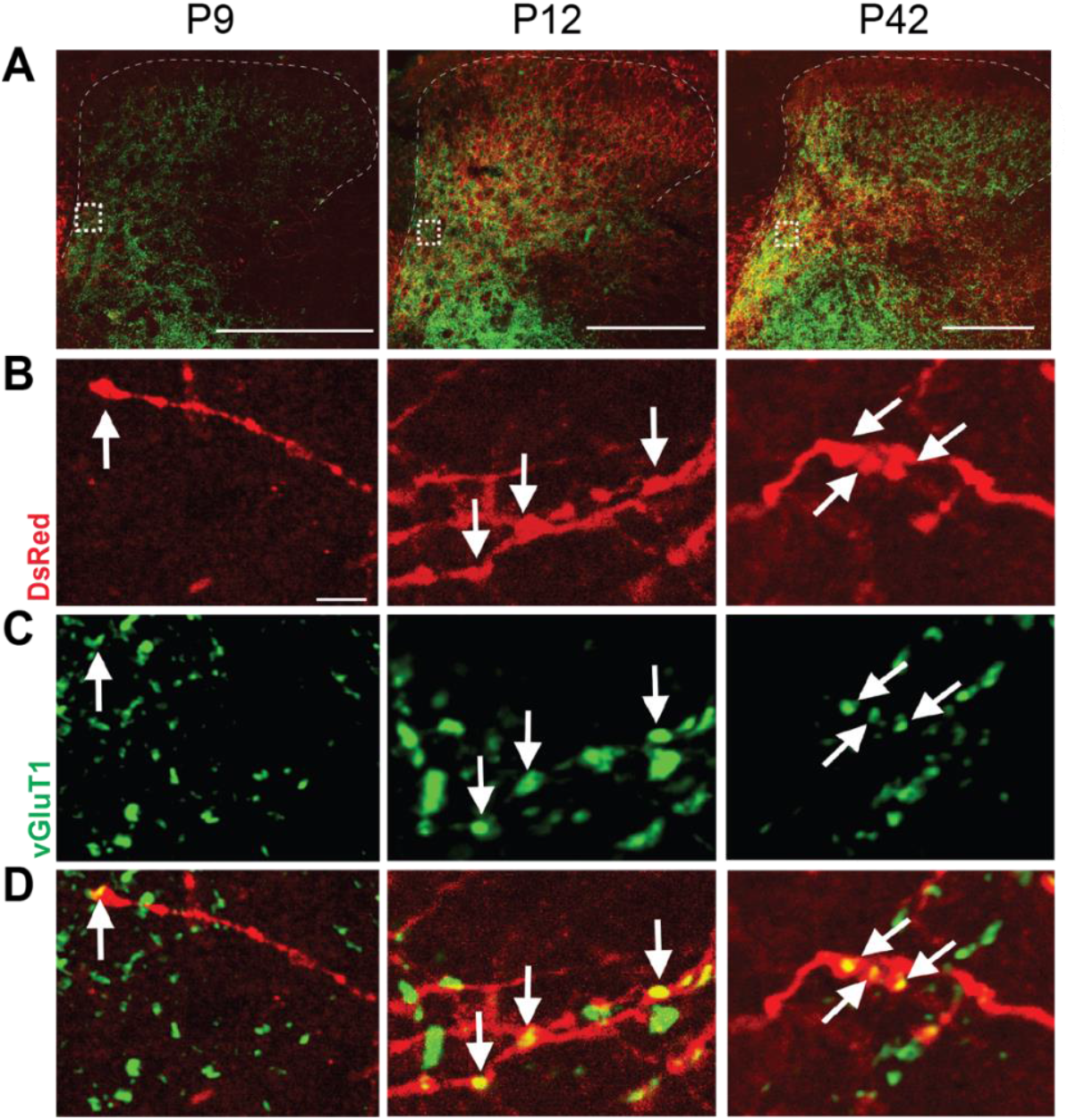
Putative S1hl-CST synaptic terminals are present in the medial deep dorsal horn from P9. **(A)** Representative lumbar spinal cord (L4) coronal sections at P9, P12, and P42 with anterogradely labelled descending S1hl-CST projections (DsRed; red) co-labelled with glutamatergic presynaptic terminals (vGluT1; green) to identify putative synaptic contacts. **(B, C, D)** High power magnification of inset from (A) showing (B) S1hl-CST fibers (DsRed), **(C)** glutamatergic terminals (vGluT1) and **(D)** merged (yellow). White arrows indicate putative excitatory synapses of S1hl-CST projections in L3-5 spinal dorsal horn, (yellow). Scale bar 500µm in (A) and 10µm in (B, C, D).

## Discussion

This study provides the first quantitative anatomical description of the postnatal maturation of S1hl-CST projections to the mouse lumbar spinal cord. By combining retrograde and anterograde tracing, we define milestones in pathway assembly: the emergence of lumbar-innervating corticospinal neurons from S1hl, the invasion and refinement of their axonal arbors within dorsal horn grey matter, and a transient laminar overextension that precedes adult-like laminar distribution.

Retrograde tracing demonstrates that S1hl corticospinal neurons projecting to lumbar dorsal horn first appear at P9 and reach adult-like numbers by P12 (Fig. 1E). This pattern aligns with classic developmental studies of the corticospinal system in rats, where early work reported the initial appearance of CST fibres at lower lumbar levels around P9 (Donatelle, 1977; Joosten, Gribnau and Dederen, 1989). By normalising FG^+^ to DAPI^+^ counts within the region of interest (Fig. 1E, Supp. Fig. 2), we show that this increase reflects a genuine increase in corticospinal neurons rather than global changes in cortical cell density, which has been reported to decline over early postnatal life due to tissue growth (Bandeira *et al*., 2009). Our analysis of anterograde labelling reveals a non-linear trajectory of S1hl-CST spinal integration into the lumbar spinal cord. Prior to P9, S1hl-CST axons are largely confined to white matter, with minimal grey matter innervation (Fig. 2C). This is followed by a period of growth defined by a rapid expansion of projections into dorsal horn grey matter between P9 and P14, reaching a peak occupancy that exceeds adult levels (Fig. 2D) and a subsequent period of refinement of axonal branching until P17. This pattern, initial exuberance followed by pruning, mirrors classic descriptions of CST development in the rat pyramidal tract and cervical spinal cord (Stanfield, O’Leary and Fricks, 1982; Curfs, Gribnau and Dederen, 1994). The temporal dissociation between cortical neuron establishment (P12) and spinal occupancy (P14-P17) suggests a structural reorganisation of axonal arbors within grey matter rather than elimination of corticospinal neurons. A similar pattern of early exuberant innervation followed by later refinement has been described for myelinated primary afferents within the dorsal horn (Beggs *et al*., 2002; Xu *et al*., 2024). In this system, refinement is activity-dependent, such that blocking spinal activity during the pruning phase disrupts mature termination patterns (Beggs *et al*., 2002). These parallels suggest that S1hl-CST maturation unfolds in an activity-dependent manner, although whether pruning is driven predominantly by intrinsic corticospinal firing, local network dynamics, or their interaction remains unresolved.

Spinal laminar targeting of CST projections has previously been suggested to act as a source of task-selectivity in the adult through laminar specific recruitment (Ueno *et al*., 2018). Our results suggest that this laminar selective circuit innervation arises late within the postnatal period. Whereas we found predominant LIII-V innervation by S1hl-CST throughout the developmental period, there was a significant fraction of superficially-projecting axons during second postnatal week (Fig. 3C-E). Transient laminar overextension has been observed in other corticospinal pathways and species, including M1-derived projections in rodents and cats (Stanfield, O’Leary and Fricks, 1982). In these cases, overextended axons are thought to participate in early circuit exploration before restraint by activity-dependent mechanisms leads to mature topography. Our findings extend this concept to the S1hl-CST: superficial laminar occupancy peaks in parallel with maximal grey matter invasion and decreases as overall territory and laminar precision consolidate. The tight temporal coupling of these processes raises the possibility that shared mechanisms such as competition with afferent inputs (Jiang *et al*., 2016) or interactions with local interneuron populations may coordinate both spatial pruning and laminar refinement.

Putative synaptic specialisation of descending cortical axons, as suggested by early colocalisation with vGluT1, emerges concurrently with axonal entry into the dorsal horn (Fig. 4). Although colocalisation does not provide functional confirmation, similar patterns have been observed in descending primary motor (M1-CST) projections in the mouse (Kamiyama, Yoshioka and Sakurai, 2006), and in kittens (Alisky, Swink and Tolbert, 1992). The presence of glutamatergic terminals from the earliest stages of grey matter engagement implies that descending somatosensory projections may influence spinal circuitry during the phase of maximal structural exuberance, positioning them as potential contributors to activity-dependent circuit refinement. Whether early superficial projections have transient functional roles or whether their removal is driven by axon-target competition, microglial engulfment, synaptic activity, or other molecular cues (Beggs *et al*., 2002; Xu *et al*., 2024) remains to be determined.

In contrast to M1-CST studies in the rat and cat, where adult-like distributions are established by the end of the second postnatal week (Donatelle, 1977; Schreyer and Jones, 1982; Curfs, Gribnau and Dederen, 1994; Martin, 2005), we find that corticospinal projections from S1hl in mouse exhibit a prolonged period of exuberant innervation and laminar overextension, with adult-like patterns reached only by the end of the third postnatal week. While direct comparison of absolute timing across species must be made cautiously, this relative delay suggests that somatosensory corticospinal pathways may retain developmental plasticity for longer than motor corticospinal projections. This extended period of refinement is consistent with the delayed maturation of dorsal horn and sensory cortical circuits (Thairu, 1971; Fitzgerald, 2005, Koch and Fitzgerald, 2013; Devonshire *et al*., 2015; Chang *et al*., 2016; Chang *et al*., 2020; Jones et al., 2022). Early in life, spinal motor output circuits support fundamental survival behaviours, including reflexive withdrawal, postural adjustments, and basic locomotor patterns (Altman and Sudarshan, 1975). Descending somatosensory cortical modulation appears to integrate subsequently, potentially refining tactile gain and enabling more context-dependent or task-selective behaviours to be mounted. In accordance with this idea, behavioural repertoires expand during our identified period of CST refinement to facilitate the transition to independent locomotion (Donatelle 1977; Altman and Sudarshan, 1975).

The transient overgrowth and subsequent pruning described here resemble developmental strategies observed across neural systems, in which early exuberant connectivity provides a substrate for activity-dependent stabilisation and circuit refinement (Shatz, 1990; Luo and O’Leary, 2005). Within the dorsal horn, transient access to both superficial and deeper laminae may allow descending cortical projections to sample multiple emerging somatosensory networks during a period of rapid spinal circuit maturation. Subsequent pruning would then consolidate corticospinal influence onto laminar territories aligned with mature sensory processing demands. In this framework, developmental overextension represents an adaptive exploratory phase rather than anatomical redundancy. Future studies combining functional readouts with targeted pathway manipulation will be necessary to determine how these structural transitions shape the emergence of adult sensorimotor integration.

## Conclusions

This work establishes a quantitative developmental framework for the somatosensory corticospinal pathway. Lumbar-projecting S1hl neurons emerge between P9 and P12, their axons invade and transiently overextend within dorsal horn grey matter, and adult-like territory and laminar specificity consolidate by the end of the third postnatal week, coincident with early synaptic specialisation.

Rather than forming with immediate target precision, the S1hl-CST appears to mature through a temporally extended phase of exuberant growth followed by selective refinement and stabilisation. This developmental trajectory places somatosensory corticospinal integration later than the maturation of gross locomotor behaviours, suggesting that descending sensory modulation is progressively layered onto pre-existing sensorimotor circuits as behavioural complexity increases.

By defining the timing and structural logic of this maturation, our findings provide a framework for future studies linking cortical activity, spinal circuit refinement, and the emergence of stable sensorimotor control. They also offer a developmental reference point for disorders in which corticospinal sensory modulation fails to consolidate appropriately.

## Author Contributions

Conceptualization, S.C.K.; Data curation, A.C., L.F., S.C.K.; Formal analysis, A.C., L.F., S.C.K. Investigation, A.C., S.C.K.; Methodology, A.C., L.F., S.C.K. Project administration, L.F., S.C.K.; Resources, L.F., S.C.K.; Validation, A.C., L.F., S.C.K.; Visualization, A.C., L.F., S.C.K.; Writing – original draft, A.C., L.F., S.C.K.; Writing – reviewing and editing, A.C., L.F., S.C.K.

## References

Abraira, V.E. and Ginty, D.D. (2013) “The sensory neurons of touch,” Neuron, 79(4), pp. 618– 639.

Alisky, J.M., Swink, T.D. and Tolbert, D.L. (1992) “The postnatal spatial and temporal development of corticospinal projections in cats,” Experimental Brain Research, 88(2), pp. 265–276. Available at: 10.1007/BF02259101.

Altman J, Sudarshan K. (1975) “Postnatal development of locomotion in the laboratory rat.” Anim Behav. 23(4), pp. 896–920. Available at: 10.1016/0003-3472(75)90114-1.

Alvarez, F.J. et al. (2004) “Vesicular glutamate transporters in the spinal cord, with special reference to sensory primary afferent synapses,” Journal of Comparative Neurology, 472(3), pp. 257–280. Available at : 10.1002/cne.20012.

Beggs, S. et al. (2002) “The postnatal reorganization of primary afferent input and dorsal horn cell receptive fields in the rat spinal cord is an activity-dependent process,” European Journal of Neuroscience, 16(7), pp. 1249–1258.

Broadhead, M.J. et al. (2020) “Nanostructural Diversity of Synapses in the Mammalian Spinal Cord,” Scientific Reports, 10(1), p. 8189. Available at: 10.1038/s41598-020-64874-9.

Catapano, A., Magavi, S. and Macklis, J. (2002) “Neuroanatomical Tracing of Neuronal Projections with Fluoro-Gold,” Methods in Molecular Biology, 198. Available at: 10.1385/1-59259-186-8:299.

Chang, P. et al. (2016) “The Development of Nociceptive Network Activity in the Somatosensory Cortex of Freely Moving Rat Pups,” Cerebral Cortex, 26(12), pp. 4513–4523. Available at: 10.1093/cercor/bhw330.

Chang, P., Fabrizi, L. and Fitzgerald, M. (2020) ‘Distinct age-dependent C fiber-driven oscillatory activity in the rat somatosensory cortex’, eNeuro, 7(5), ENEURO.0036-20.2020. doi: 10.1523/ENEURO.0036-20.2020.

Curfs, M.H.J.M., Gribnau, A.A.M. and Dederen, P.J.W.C. (1994) “Selective elimination of transient corticospinal projections in the rat cervical spinal cord gray matter,” Developmental Brain Research, 78(2), pp. 182–190. Available at: 10.1016/0165-3806(94)90025-6.

Devonshire, I.M., Greenspon, C.M. and Hathway, G.J. (2015) ‘Developmental alterations in noxious-evoked EEG activity recorded from rat primary somatosensory cortex’, Neuroscience, 305, pp. 343–350. doi: 10.1016/j.neuroscience.2015.08.004.

Donatelle, J.M. (1977) “Growth of the corticospinal tract and the development of placing reactions in the postnatal rat,” Journal of Comparative Neurology, 175(2), pp. 207–231. Available at: 10.1002/cne.901750205.

Fitzgerald, M. (2005) “The development of nociceptive circuits,” Nature Reviews Neuroscience, 6(7), pp. 507–520. Available at : 10.1038/nrn1701.

Frezel, N. et al. (2020) “In-Depth Characterization of Layer 5 Output Neurons of the Primary Somatosensory Cortex Innervating the Mouse Dorsal Spinal Cord,” Cerebral Cortex Communications, 1(1). Available at: 10.1093/texcom/tgaa052.

Gribnau, A.A.M. et al. (1986) “On the development of the pyramidal tract in the rat,” Anatomy and Embryology, 175(1), pp. 101–110. Available at : 10.1007/BF00315460.

Guan, X. et al. (2025) “Neuropathic Pain-Like Responses in a Chronic CNS Injury Model Are Mediated by Corticospinal-Targeted Spinal Interneurons,” The Journal of Neuroscience, 45(29), p. e1264242025. Available at: 10.1523/JNEUROSCI.1264-24.2025.

Jiang Y. Q., Zaaimi B., Martin J. H. (2016) “Competition with Primary Sensory Afferents Drives Remodeling of Corticospinal Axons in Mature Spinal Motor Circuits”. The Journal of Neuroscience, 6;36(1) pp.:193–203. Available at: 10.1523/JNEUROSCI.3441-15.2016.

Jones, L., Verriotis, M., Cooper, R.J., Laudiano-Dray, M.P., Rupawala, M., Meek, J., Fabrizi, L. and Fitzgerald, M. (2022) ‘Widespread nociceptive maps in the human neonatal somatosensory cortex’, eLife, 11, e71655. doi: 10.7554/eLife.71655.

Joosten, E.A.J., Gribnau, A.A.M. and Dederen, P.J.W.C. (1989) “Postnatal development of the corticospinal tract in the rat,” Anatomy and Embryology, 179(5), pp. 449–456. Available at: 10.1007/BF00319587.

Kamiyama, T., Yoshioka, N. and Sakurai, M. (2006) “Synapse Elimination in the Corticospinal Projection During the Early Postnatal Period,” Journal of Neurophysiology, 95(4), pp. 2304– 2313. Available at: 10.1152/jn.00295.2005.

Koch, S.C. and Fitzgerald, M. (2013) “Activity-dependent development of tactile and nociceptive spinal cord circuits,” Annals of the New York Academy of Sciences, 1279(1), pp. 97–102. Available at: 10.1111/nyas.12033.

Lemon, R.N. and Griffiths, J. (2005) “Comparing the function of the corticospinal system in different species: Organizational differences for motor specialization?” Muscle & Nerve, 32(3), pp. 261–279. Available at : 10.1002/mus.20333.

Liu, Y. et al. (2018) “Touch and tactile neuropathic pain sensitivity are set by corticospinal projections,” Nature, 561(7724), pp. 547–550. Available at: 10.1038/s41586-018-0515-2.

Luo, L. and O’Leary, D.D.M. (2005) “Axon retraction and degeneration in development and disease” Annual Review of Neuroscience, 28, pp. 127–156. Available at: 10.1146/annurev.neuro.28.061604.135632

Malmberg, A.B. et al. (1997) “Preserved Acute Pain and Reduced Neuropathic Pain in Mice Lacking PKCγ,” Science, 278(5336), pp. 279–283. Available at: 10.1126/science.278.5336.279.

Martin, J.H. (2005) “The Corticospinal System: From Development to Motor Control,” The Neuroscientist, 11(2), pp. 161–173. Available at : 10.1177/1073858404270843.

Moreno-Lopez, Y. et al. (2021) “The corticospinal tract primarily modulates sensory inputs in the mouse lumbar cord,” eLife. Edited by M. Thoby-Brisson et al., 10, p. e65304. Available at: 10.7554/eLife.65304.

Schreyer, D.J. and Jones, E.G. (1982) “Growth and target finding by axons of the corticospinal tract in prenatal and postnatal rats,” Neuroscience, 7(8), pp. 1837–1853. Available at: 10.1016/0306-4522(82)90001-X.

Shatz, C.J. (1990) “Impulse activity and the patterning of connections during CNS development” Neuron, 5(6), pp. 745–756. Available at: 10.1016/0896-6273(90)90333-B.

Stanfield, B.B., O’Leary, D.D. and Fricks, C. (1982) “Selective collateral elimination in early postnatal development restricts cortical distribution of rat pyramidal tract neurones.,” Nature, 298(5872), pp. 371–3. Available at: 10.1038/298371a0.

Thairu, B.K. (1971) “Post-natal changes in the somaesthetic evoked potentials in the albino rat”, Nature New Biology, 231, pp. 30–31. doi: 10.1038/newbio231030a0.

Ueno, M. et al. (2018) “Corticospinal Circuits from the Sensory and Motor Cortices Differentially Regulate Skilled Movements through Distinct Spinal Interneurons,” Cell Reports, 2018/05/03, 23(5), pp. 1286-1300.e7. Available at : 10.1016/j.celrep.2018.03.137.

Varoqui, H. et al. (2002) “Identification of the Differentiation-Associated Na ^+^ /P _I_ Transporter as a Novel Vesicular Glutamate Transporter Expressed in a Distinct Set of Glutamatergic Synapses,” The Journal of Neuroscience, 22(1), pp. 142–155. Available at : 10.1523/JNEUROSCI.22-01-00142.2002.

Wang, G.-H. et al. (2025) “Descending projection neurons in the primary sensorimotor cortex regulate neuropathic pain and locomotion in mice,” Nature Communications, 16(1), p. 5918. Available at : 10.1038/s41467-025-61164-8.

Yamawaki, N. et al. (2014) “A genuine layer 4 in motor cortex with prototypical synaptic circuit connectivity” eLife 3:e05422. Available at : https://elifesciences.org/articles/05422

Xu, Y., et al. (2024) ‘Microglial refinement of A-fiber projections in the postnatal spinal cord dorsal horn is required for normal maturation of dynamic touch’, Journal of Neuroscience, 44(2), e1354232023. doi: 10.1523/JNEUROSCI.1354-23.2023.

