## Supplemental Figures for "Postnatal development of somatosensory corticospinal projections in the mouse lumbar spinal cord"

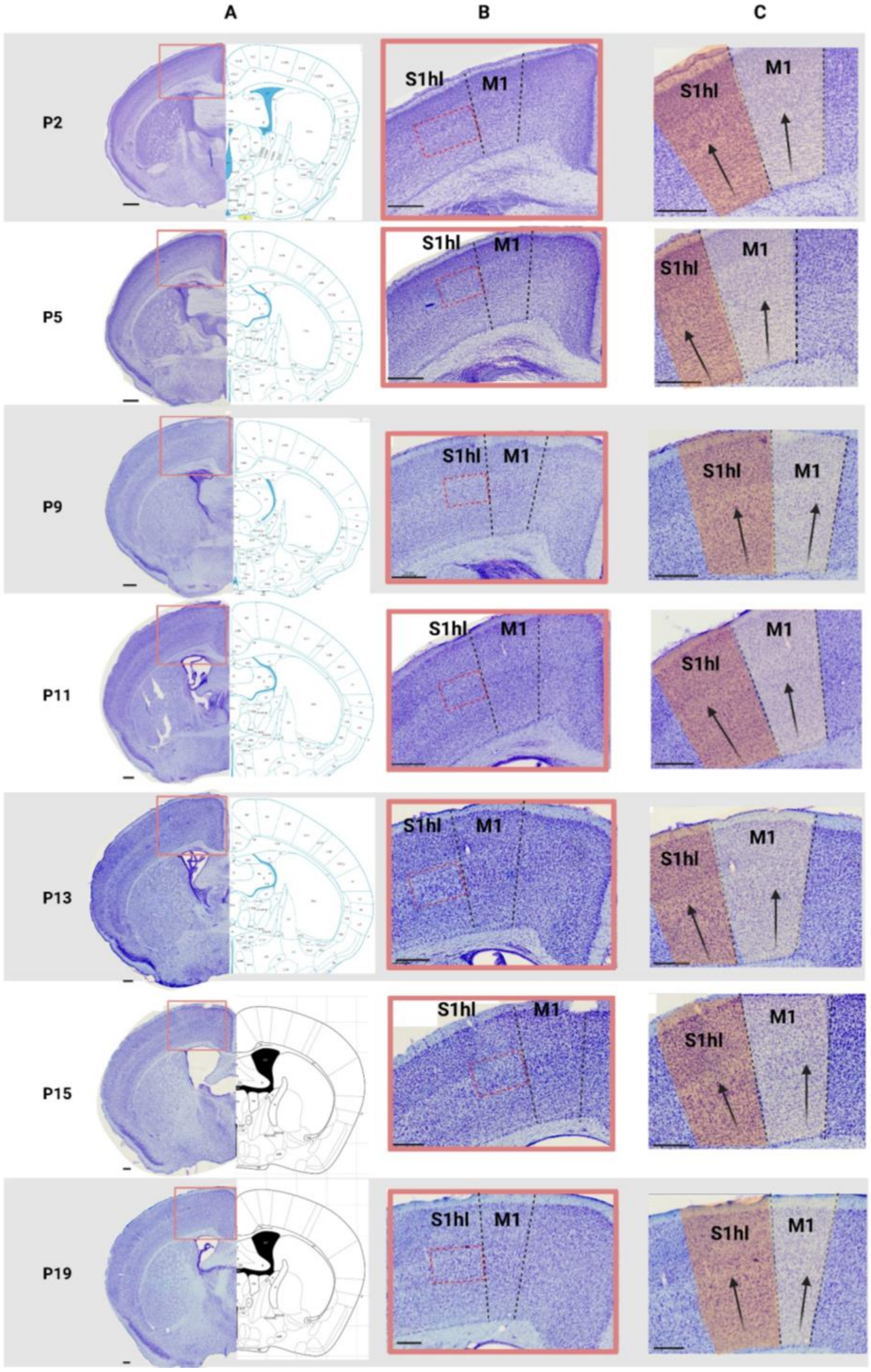


**Supp. Fig. 1 Representative Nissl-stained coronal sections of the brain at P2, P5, P9, P11, P13, P15 and P19 to define S1hl boundaries. A)** (left) Nissl-stained coronal hemisections of 7 postnatal ages and (right) corresponding reference stereotaxic atlas: P0 atlas for P2, P6 atlas for P5, P9, P11, and P13, and adult atlas for P15 and P19. **B)** High magnification of highlighted areas in A) showing delineation of M1 and S1hl (black dotted lines); dashed red boxes highlight layer 5 of S1hl. **C)** Higher magnification of cortical area illustrating M1 (yellow) and S1hl (orange) delineation based on two complementary features: (1) cellular orientation between cortical columns S1/M1 (white arrows), and (2) L4/L5 thickness and cellular packing. Scale bar: 200μm.

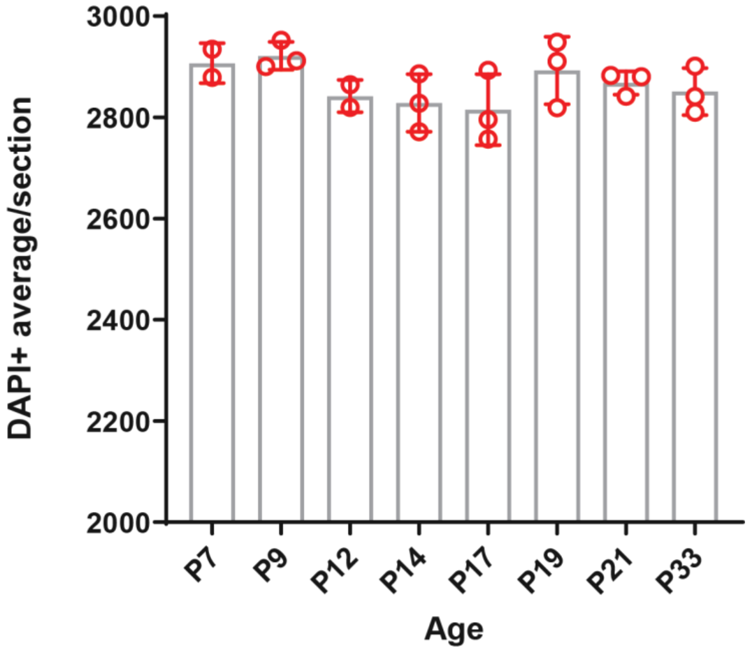


**Supp. Fig. 2. Cortical DAPI⁺ cell counts do not differ between ages:** A Quantification of average DAPI⁺ cells within S1hl L5 across postnatal development. Group comparisons for DAPI counts were performed using one-way ANOVA with a post-hoc Tukey's multiple comparisons test ($p<0.05$) No significant difference was observed between ages. Results presented as mean ± SD. Scale bar 100μm.
